# A default silencing mechanism restrains stress-induced genes in *C. elegans*

**DOI:** 10.1101/2025.10.22.683740

**Authors:** Orkan Ilbay, Alejandro Rodriguez Gama, Daniel F Jarosz, Richard I. Morimoto, Andrew Fire

## Abstract

Inducible gene expression programs require that target genes remain silent until the proper activation signal is received, a hallmark of stress-response pathways. This quiescence is typically assumed to be the default state of stress-inducible genes, maintained without active cellular intervention. Using a forward genetic screen for constitutive activation of inducible heat shock proteins (iHSPs) in *C. elegans*, we found that the multi-zinc-finger protein ZNF-236 is essential for maintaining iHSP quiescence under normal conditions. Loss of *znf-236* causes constitutive iHSP expression throughout the genome, affecting both endogenous iHSP loci and iHSP sequences inserted at dispersed chromosomal sites. However, the effect is also chromosomal context-dependent: robustly heat-responsive iHSP transgenes integrated into the ribosomal DNA locus or extrachromosomal arrays are unaffected by *znf-236* loss. This differential responsiveness suggests iHSP induction in *znf-236* mutants results from a shift in genome organization, rather than from accumulated denatured proteins or engagement of the canonical heat shock response. Our findings demonstrate the existence of a potent ZNF-236-dependent default silencing mechanism that broadly restrains iHSP genes across the genome and helps ensure appropriate iHSP quiescence even at ectopic chromosomal locations. Contrary to prior assumptions, this suggests that quiescence of stress-inducible genes reflects an actively maintained genomic state rather than merely a passive absence of expression.

**Graphical abstract:** 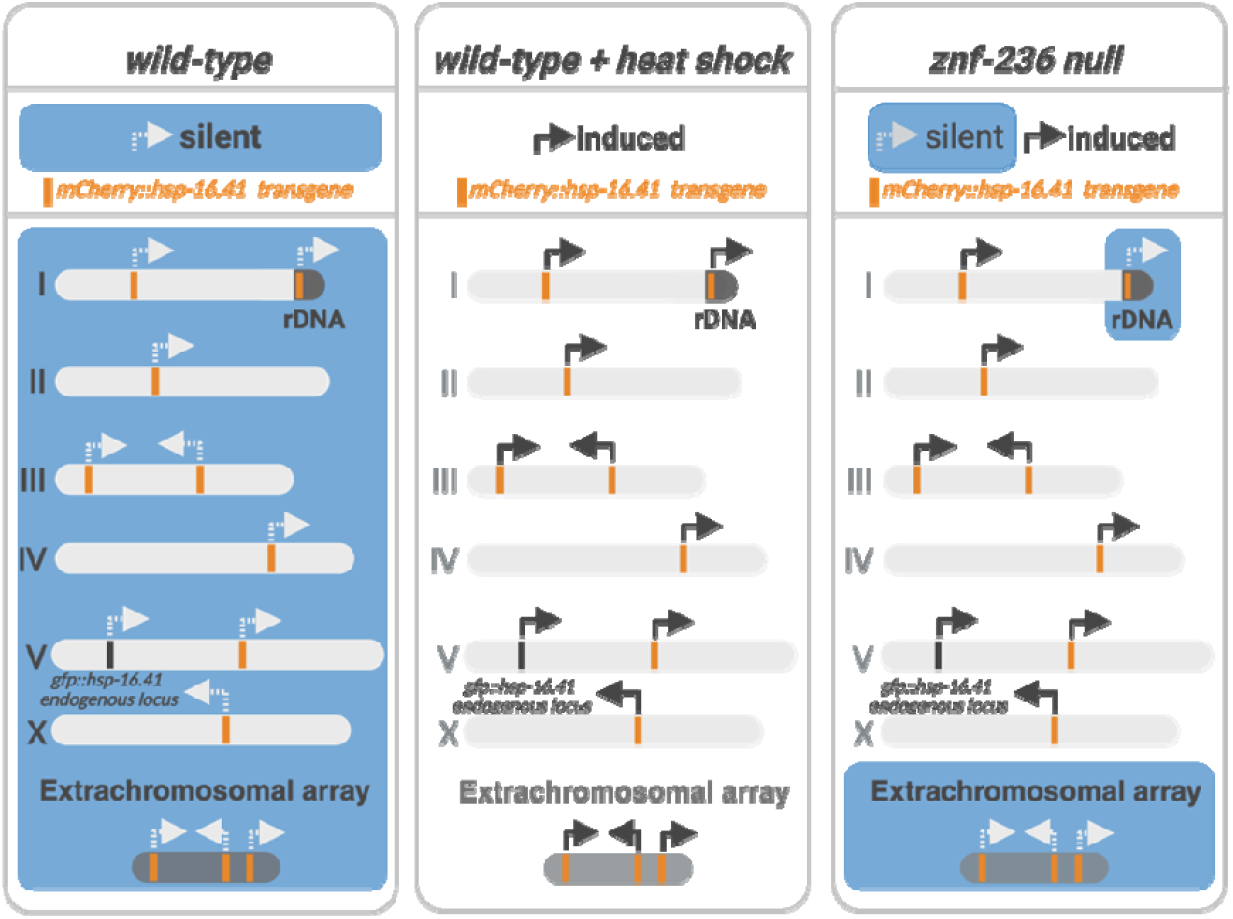

## INTRODUCTION

Despite constantly changing environmental conditions, biological systems are remarkably successful at maintaining normal cellular functions. Even under severe stress that damages cellular biomolecules, specialized programs activated by stress-response pathways mitigate damage, restore homeostasis, and enhance survival. The defining feature of these programs is the rapid induction of stress- or damage-responsive genes that are normally silent. The baseline silence under nonstress conditions is typically assumed to be a passive default rather than an actively enforced state.

Elevated temperatures pose a universal challenge to living cells, particularly affecting proteins, which are highly sensitive to heat stress. At high temperatures, cellular proteins can denature, losing their functional three-dimensional structures and exposing their hydrophobic, aggregation-prone core regions. Denatured proteins activate the widely conserved transcription factor heat-shock factor 1 (HSF-1)^1,2^, which mediates the heat shock response (HSR)^3–5^. HSF-1 promotes the expression of inducible heat-shock proteins (iHSPs), molecular chaperones that help mitigate protein damage and restore homeostasis once the conditions return to normal.

Like other stress response pathways, the heat-shock response is thought to have evolved to address transient insults, functioning most effectively when activated in proportion to the intensity and duration of stress^6^. This regulation is achieved by transient activation of HSF-1. Although HSF-1 is constitutively expressed, it remains inactive under normal conditions. Its activation and subsequent deactivation during and after stress involve multiple post-translational steps^7–9^, with chaperone binding playing a central role in both processes^2,10,11^. Under non-stress conditions, constitutively expressed chaperones bind and inhibit HSF-1, while during stress, denatured proteins engage these chaperones, freeing HSF-1 to initiate HSR. As the response progresses, molecular chaperones are replenished and rebind to HSF-1, restoring its inactive state and attenuating the HSR. Thus, the HSR is driven by a finely tuned cycle of HSF-1 release and re-inactivation, ensuring an adaptive but self-limiting response to proteotoxic stress.

Constitutive HSF-1 activation or excess HSP expression can be detrimental. For example, in fruit flies, ectopic expression of a single inducible heat-shock protein alone is harmful^12^. Moreover, persistent HSF-1 activation and HSP overexpression, frequently observed in various cancers, contribute to malignant transformation and poor prognosis^13–16^. Therefore, maintaining the quiescence of the stress-inducible HSPs (iHSPs) is crucial for cellular and organismal health. While this quiescence can be largely attributed to chaperone-mediated inhibition of HSF-1, additional mechanisms that prevent the expression of iHSPs or other stress-induced genes under normal conditions --as well as factors leading to constitutive HSP expression--remain unknown.

*Caenorhabditis elegans (C. elegans)* is a free-living soil nematode that naturally encounters fluctuating environmental conditions, making it an ideal organism for studying stress responses *in vivo*. It has emerged as a central model for dissecting the metazoan heat-shock response (HSR), owing to its genetic tractability, transparent body, and well-annotated genome^4^. Like other systems, in *C. elegans*, the heat-shock factor HSF-1 is essential and plays a central role in the HSR by recognizing conserved heat-shock elements (HSEs) in the promoters of inducible heat-shock protein genes^17,18^. The fundamental mechanisms of HSF-1 activation—including trimerization, DNA binding, and transcriptional activation—are highly conserved from worms to mammals^19^.

Beyond these core molecular features, *C. elegans* offers unique advantages for studying heat-shock biology in the context of a whole organism, facilitating discoveries about HSF-1’s role in longevity and development^20–25^, as well as the non-cell autonomous, organism-wide regulation of the heat-shock response by the nervous system^26–28^.

In this study, we identified the multi-zinc-finger protein ZNF-236 as a key regulator of expression of inducible heat-shock proteins and a subset of stress-induced prion-like proteins in *C. elegans*. The capacity for constitutive activation of stress-induced genes in the *znf-236* mutants is evidently genome-wide, with two observed exceptions: rDNA and extrachromosomal arrays. Our findings are consistent with a ZNF-236-dependent default silencing mechanism that broadly restrains iHSP genes across the genome. This potent mechanism ensures quiescence of iHSP genes even when inserted at ectopic chromosomal locations and demonstrates that quiescence of stress-inducible genes can be an actively maintained genomic state rather than merely a passive absence of expression.

## RESULTS

### A forward genetic screen identifies *znf-236* as a negative regulator of the heat shock gene expression

Although *C. elegans* is typically cultured at temperatures ranging from 15°C to 25°C, the species can survive short exposures to higher temperatures--such as 35°C--for a few hours. During such stress, the heat shock response (HSR), mediated by the transcription factor HSF-1, is activated to mitigate cellular damage and restore homeostasis where possible.

We hypothesized that while prolonged exposure to HSR-inducing temperatures is lethal, constitutive activation of the heat shock response (HSR) due to mutations in the genetic pathways upstream of *hsf-1* might be tolerated in *C. elegans*. To identify and study such genetic events, we required a sensitive and tractable HSR reporter that would be compatible with forward mutagenesis screens—specifically, one that allows the distinction between cis-acting mutations (e.g. in the promoter of the reporter) and trans-acting regulatory factors. To this end, we generated a single-copy reporter by tagging *hsp-16*.*41*, a well-characterized HSF-1 target that encodes a small heat-shock protein (orthologous of human crystallin alpha A), with a fluorescent protein at its native genomic locus (**Figure 1A**).

**Figure 1.**
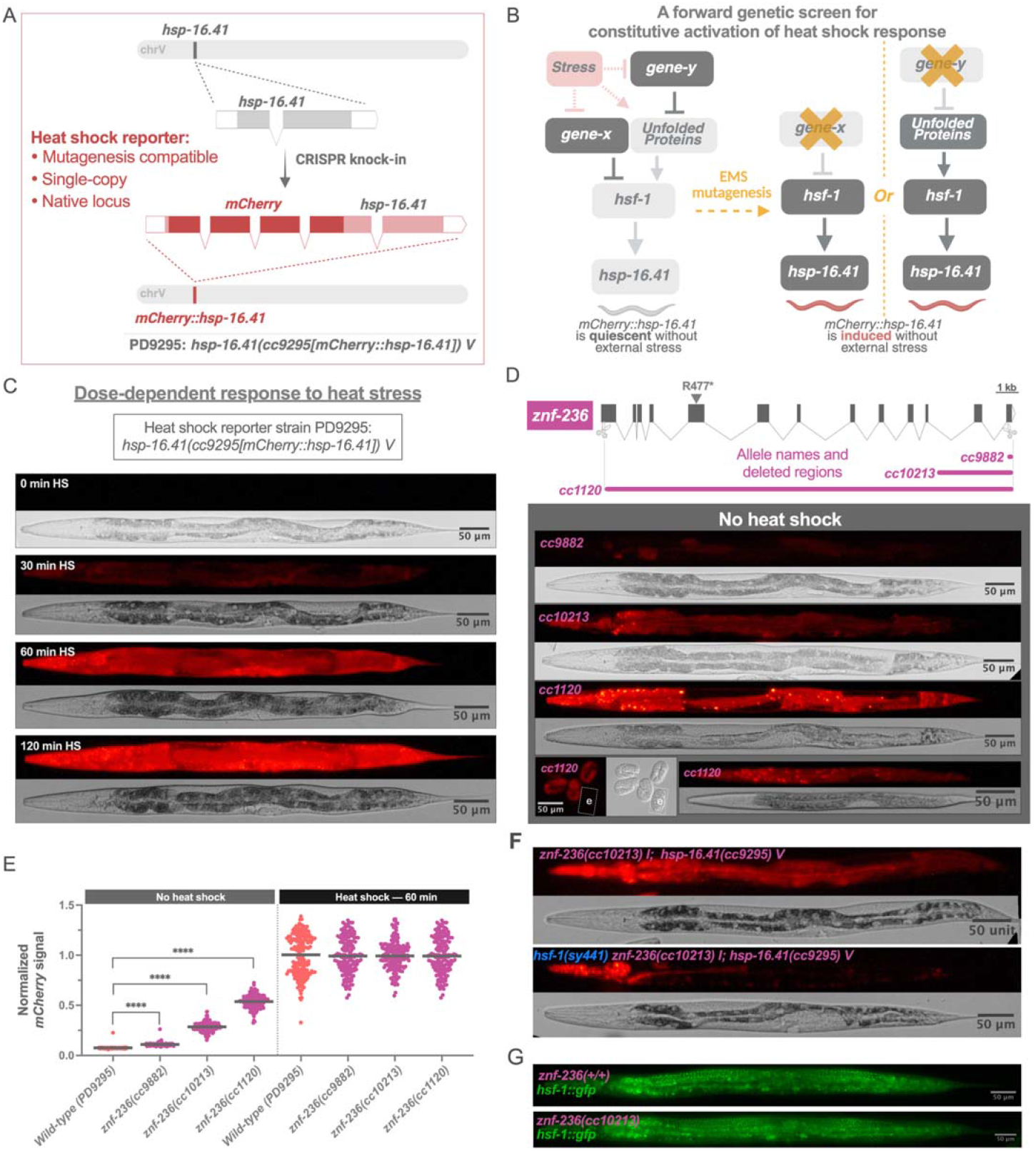
A forward genetic screen identifies *znf-236* as a negative regulator of heat shock gene expression. (A) Schematic of the *hsp-16*.*41* locus tagged with *mCherry* to generate the heat shock reporter strain PD9295. (B) Rationale behind the forward genetic screen: Genetic factors integrating stress signals into the heat shock response by suppressing HSF-1, either directly (gene-x) or indirectly by preventing proteotoxic stress (gene-y), may normally prevent *hsp-16*.*41* induction. Loss-of-function mutations in such genes are predicted to cause constitutive expression of the *mCherry::hsp-16*.*41* reporter. (C) Representative micrographs showing *mCherry::hsp-16*.*41* expression under normal conditions (0 min heat shock), and after heat shock (HS) at 35°C for 30, 60, or 120 minutes, followed by 5 hours of recovery to allow response maturation. Identical microscopy settings were used for all condition. (D) Gene structure of *znf-236*, showing exon-intron structure, CRISPR-generated deletion alleles, and the region disrupted by each allele. The R477* allele was identified in the EMS screen. Micrographs show constitutive *mCherry::hsp-16*.*41* induction in each *znf-236* mutant in the absence of heat shock. In the *cc1120* panel, ‘e’ marks an early embryo lacking induction; bottom right image shows induction in an L2 larva. All images in (C) and (D) were acquired under the same microscopy settings. (E) *mCherry::hsp-16*.*41* expression signal measured by high-throughput imaging of the indicated *znf-236* alleles. Under no heat shock, mutant alleles display significant induction of *mCherry::hsp-16*.*41*. After a 35 °C for 60 minutes followed by 5 hours recovery, mutant alleles display induction of mCherry *mCherry::hsp-16*.*41* of around 50% for cc1120, 75% for cc10213 and 90% for cc9882. Each group contains at least 60 worms. **** P-val <0.001. (F) *mCherry::hsp-16*.*41* expression in *znf-236(cc10213)* mutants carrying a partial loss-of-function mutation of *hsf-1*, showing reduced reporter induction. (G) HSF-1::GFP expression in wild-type and a *znf-235(cc10216)* worm, showing comparable levels.

In addition to its compatibility with mutagenesis, a key advantage of our reporter (**Figure 1A**) is its ability to sense, respond to, and thus reveal regulatory input from both trans-acting factors (e.g., upstream gene products) and a broad range of cis-acting elements, including both proximal and distal regulatory sequences that control the endogenous *hsp-16*.*41* locus.

The heat shock response (HSR), or the iHSPs, can be activated not only by elevated temperatures but also by other stressors that disrupt proteostasis, including oxidative stress, heavy metals, and pathogens^5^. Denatured proteins alone are sufficient to trigger HSP induction^1^, and are thought to do so by sequestering constitutively expressed HSPs, thereby freeing and activating HSF1^1,29^. However, whether protein unfolding or misfolding is a necessary component of HSF-1 activation under various sources of stress remains unclear.

We hypothesized that there might be genetic factors acting upstream of *hsf-1* that either indirectly suppress HSF-1 activation by promoting protein homeostasis^30^ --and thus preventing protein unfolding-- or directly sensing proteotoxic or other stress signals and integrating them into the HSR independently of unfolded proteins (**Figure 1B**). In either case, loss-of-function mutations in these upstream factors would be expected to activate the HSR even in the absence of external stresses (**Figure 1B**), a response that could be detected by our HSR reporter **(Figure 1A)**.

First, we confirmed that the tagged *hsp-16*.*41* reporter remains quiescent at normal culture temperatures (below 25°C) and is robustly activated upon exposure to elevated temperatures (**Figure 1C**). The strength of the observed induction correlates with the duration of the heat stress (**Figure 1C**).

Next, we mutagenized^31^ late L4 stage worms carrying the HSR reporter (**Figure 1A**: Strain designation PD9295), and screened F1 and F2 progeny for visible mCherry signal under a dissecting microscope--indicative of HSR induction without external stress. Consistent with previous reports on the detrimental effects of constitutive HSR activation, the majority of ~800 mutagenized animals showing ectopic mCherry expression either failed to reach adulthood or were sterile as adults. Nevertheless, we established several stable lines exhibiting persistent mCherry expression (**Supplemental figure 1**). In some of these lines, mCherry was expressed broadly across nearly all tissues, while in others, expression was restricted to a single tissue, such as body wall muscle cells or intestine (**Supplemental figure 1**)—suggesting the tissue-specific engagement of proteostasis or stress signaling pathways, as previously reported^30^.

We focused on one mutant line that showed widespread *hsp-16*.*41* induction in multiple tissues, resembling the induction pattern observed following heat shock. We performed whole genome sequencing, and identified a premature stop codon, R477* in a candidate causative gene *znf-236* (**Figure 1D)**. Using CRISPR/Cas9 genome editing, we generated multiple *znf-236* alleles, including a ~17 kb deletion of the entire *znf-236* open reading frame, *znf-236(cc1120) (***Figure 1D**). *znf-236* mutations led to constitutive activation of *hsp-16*.*41*, confirming R477* as the causative mutation—and indicating that no compensatory mutation is required in the background to support viability (**Figure 1D**).

The strength of *hsp-16*.*41* induction without external stress varied in different *znf-236* mutants, ranging from milder induction in *cc9882 and cc10213* (partial loss of function alleles), to the strongest induction in the *cc1120* (loss of function allele) (**Figure 1D**). In the *cc1120* allele, *hsp-16*.*41* induction can be detected as early as late-stage embryos, across multiple cell types, including neurons, pharyngeal, intestinal, muscle, and coelomocyte cells-–without evident expression in the germline or hypodermis. This induction persists throughout larval development into adulthood -- with full penetrance in adults (**Figure 1D, E**). Despite this continuous HSR-like activity, all mutants remain responsive to heat stress: *mCherry::hsp-16*.*41* expression can be further induced 50 percent more in the *cc1120*, allele which already displays basal expression higher than wild-type worms after heat shock (**Figure 1E**).

ZNF-236, the protein product of the *znf-236* gene, is a zinc-finger protein containing multiple C2H2 zinc-finger domains--22 in total, according to InterProScan. ZNF-236 has not previously been ascribed a biological function. Both the human and *C. elegans* genomes encode hundreds of C2H2 zinc-finger proteins, some of which are presumed orthologs across species^32^. However, due to the widespread conservation of the stereotypical zinc-finger domain signature and the rapid evolution of the non-zinc-finger regions^32^, a single human ortholog of ZNF-236 was not evident. Nevertheless, human zinc-finger proteins ZNF91, ZNF208, and ZNF845 share the highest sequence homology with ZNF-236.

We found that the constitutive induction of *hsp-16*.*41* in *znf-236* mutants is largely HSF-1 dependent, suggesting that *znf-236* acts through HSF-1 in regulating the heat shock response (**Figure 1F)**. HSF-1 levels in *znf-236* mutants appear to be comparable to non-heat-shocked wild-type, arguing against ZNF-236 regulating HSP targets through upregulation of HSF-1 abundance (**Figure 1G**). ZNF-236’s capacity to regulate HSF-1’s activity or *hsp-16*.*41* expression appears to be cell-autonomous. In chimeric worms—constructed as described before^33,34^—where the *znf-236* homozygous mutant allele is present in only one cell lineage (AB or P) and the converse lineage (P or AB) contains the homozygous wildtype *znf-236* allele, hsp*-16*.*41* expression is exclusively observed in the mutant lineage (**Supplemental Figure 2**).

**Figure 2.**
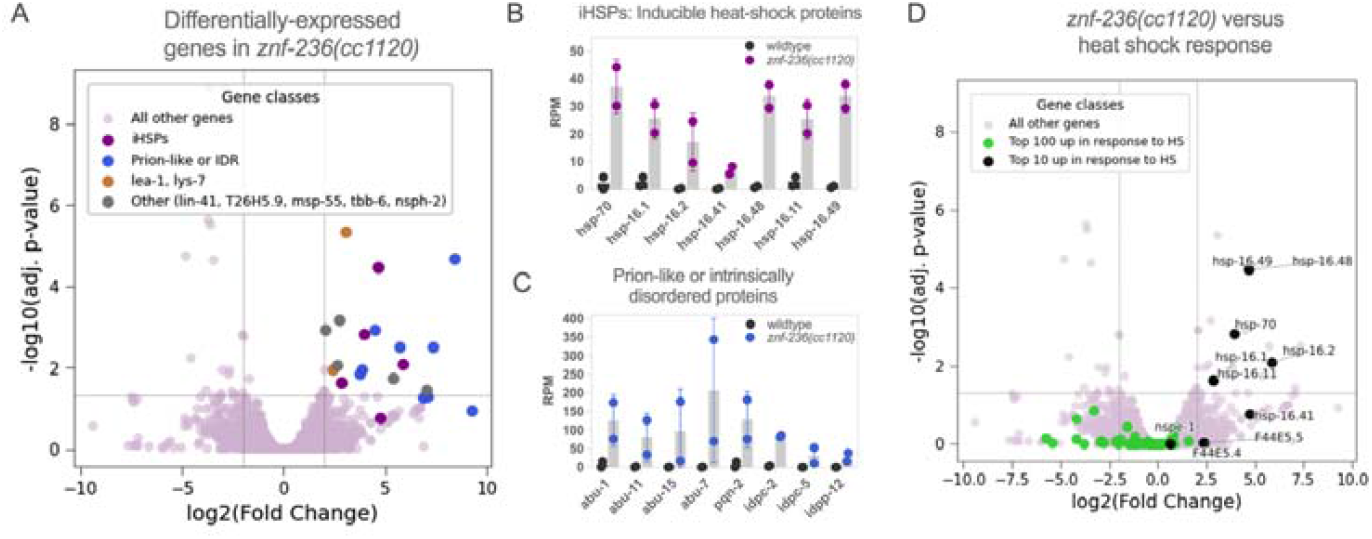
Inducible heat shock proteins (iHSPs) and a set of stress-associated prion-like or disordered proteins apparently upregulated in *znf-236* mutants. (A) Differential gene expression analysis (DESeq2) comparing L4-stage *znf-236(cc1120)* mutants to stage-matched wild-type animals, shown as a volcano plot. Upregulated gene classes are labelled. (B) Expression levels (reads per million, RPM) of inducible heat shock proteins (iHSPs) in wild-type and *znf-236(cc1120)* mutants. (C) Expression levels (RPM) of upregulated prion-like or intrinsically disordered proteins in wild-type and *znf-236(cc1120)* mutants. (D) Genes previously reported as heat shock (HS) induced (Brunquell *et al*., 2026, [36]) are highlighted on the volcano plot: black dots indicate genes ranked in the top 10, and lime green dots indicate genes ranked in top 100. The primary overlap between heat shock response and response to *znf-236* inactivation consists of iHSPs.

### Inducible heat shock proteins (iHSPs) and a set of prion-like or disordered proteins show upregulation in *znf-236* mutants

ZNF-236 prevents the inappropriate (uninduced) expression of *hsp-16*.*41*, a primary target of HSF-1 that is rapidly activated upon heat shock^35^. Because ZNF-236 also acts at least in part through HSF-1 **(Figure 1F)**, possible models explaining these observations range from ZNF-236 functioning as a direct repressor of *hsp-16*.*41* and perhaps other genes, to it acting as a broader suppressor of protein misfolding and, consequently, the full-scale heat shock response (HSR) mediated by HSF-1. To better understand the role of ZNF-236 within the HSR, we asked which other genes are differentially expressed in the *znf-236* mutants and how these changes compare to a canonical HSR. To address this, we quantified transcriptomic changes in *znf-236(cc1120)* mutants relative to stage-matched wild-type animals using high-throughput RNA sequencing.

We found that several genes are differentially expressed in *znf-236(cc1120)* mutants, including other inducible heat-shock proteins (iHSPs), such as members of the *hsp-16* family and *hsp-70* (**Figure 2A, B**), consistent with a broader engagement of the HSR pathway. Although these iHSPs are primary HSF-1 targets, not all previously identified HSF-1 targets^35,36^ are differentially expressed in *znf-236* mutants. Likewise, not all differentially expressed genes in *znf-236(cc1120)* mutant background are known HSF-1 targets (**Figure 2D)**.

Examining the differential expression in detail, the seven upregulated iHSPs in *znf-236* mutants are among the top ten most highly induced genes upon heat shock^36^ (**Figure 2D**). Five of these seven iHSPs are also among the most rapidly induced genes in response to heat shock in *C. elegans*^35^. These findings suggest that *znf-236* inactivation does not lead to full-scale/full-body activation of the HSR, but it rather leads to the selective activation of a specific subset of inducible heat shock proteins that are also rapidly and strongly induced in an *hsf-1*-dependent manner upon heat shock^35^.

In addition to iHSPs, we observed upregulation of a distinct set of prion-like genes that have previously been included in sets of stress-responsive genes (**Figure 2A, C**), including *abu-1, abu-7, abu-11, abu-15* (members of the abu family: named based on their apparent activation in blocked unfolded protein response (UPR)^37^ —which are also referred to as noncanonical UPR genes^38^. Although consistent here with a stress response induction, we note that an overlapping set of *abu-* genes (12 of 15) and other prion-like proteins have also been shown to be upregulated in pharyngeal muscles in molting animals with no defined external stress^39^, raising the possibility that elevated abu levels in *znf-236*(*cc1120)* animals could reflect features of spatial or timing control in these mutants. Of note, only a small subset of (4 or 15) abu genes are upregulated in *znf-236(cc1120)* mutants. Other upregulated genes include *pqn-2, idpc-2, idpc-5, idpp-12*, which encode prion-like or disordered proteins that are also typically quiescent under non-stress conditions (**Figure 2C)**.

In addition to iHSPs and Abu/prion-like proteins, we also observed increased expression of two other stress-related genes: *lys-7*, a pathogen-inducible lysozyme^40^, and *lea-1*, which is induced by desiccation stress^41^ (**Figure 2A**).

In summary, heat shock and loss of *znf-236* shared activation of a set of inducible heat shock proteins (iHSPs), but differ markedly beyond this class. These findings are consistent with a model in which *znf-236* is required to maintain the quiescence of stress-induced genes in two major categories: iHSPs and prion-like proteins.

### *znf-236* mutants exhibit increased thermotolerance and maintain constitutive *hsp-16*.*41* expression during aging

To assess how the altered gene expression program in *znf-236* affects physiology, especially in the context of responding to stress, we measured thermotolerance (the ability to withstand heat stress).

Employing a heat shock regimen of 6 hours at 34°C, we found that the constitutive hsp-expressing *znf-236* mutant worms displayed a significant increase in survival in this heat shock challenge (**Figure 3A**), indicating that *znf-236* mutation confers thermotolerance. The effect correlated with the degree of *hsp-16*.*41* induction, ranging from almost no change in the *cc9882* allele to substantial improvement in the *cc1120* allele.

**Figure 3.**
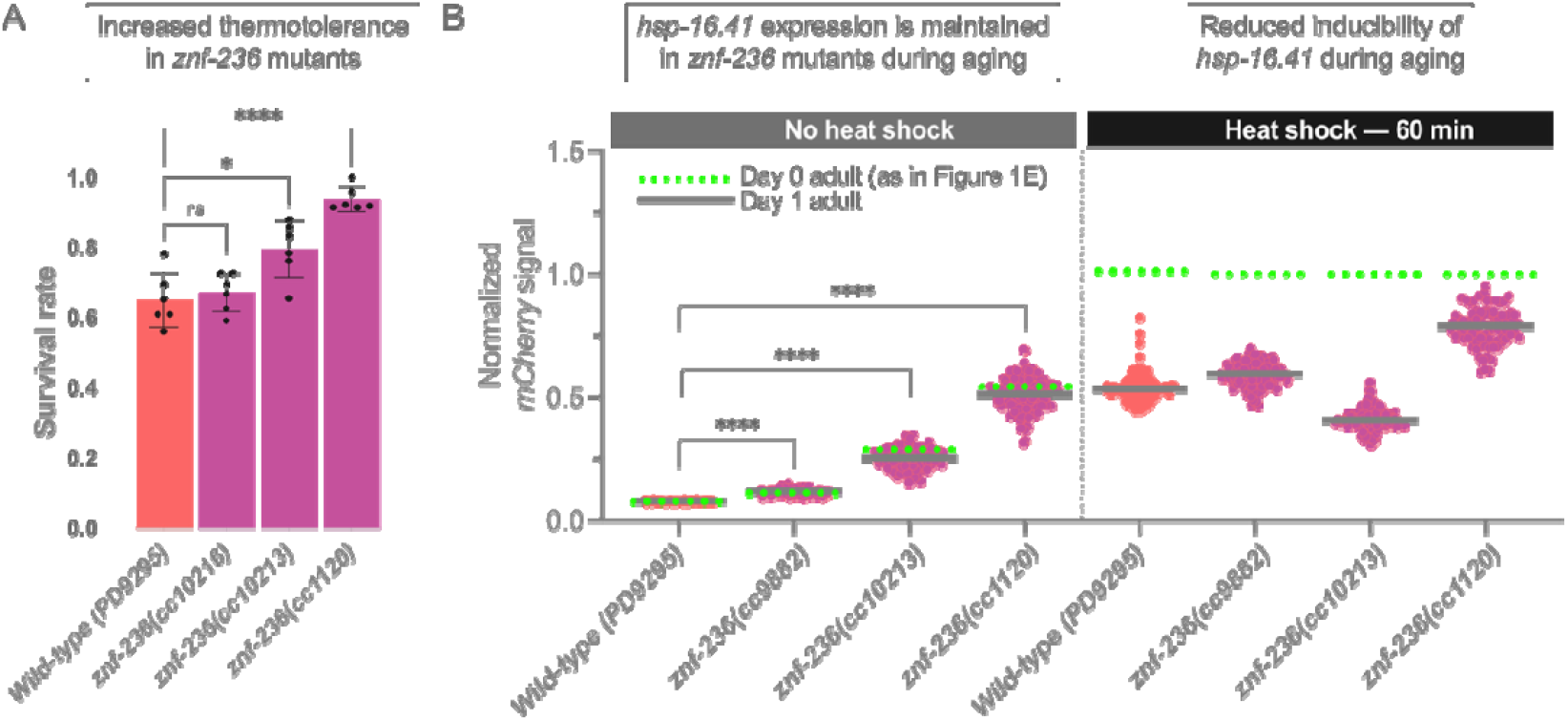
*znf-236* mutants exhibit thermotolerance and maintain *hsp-16*.*41* induction during aging. (A) Thermotolerance assay of animals exposed to 34°C for 6 hours and scored 24 hours after. n=6 with sample size of 25 animals. (B) High throughput imaging of animals expressing *mCherry::hsp-16*.*41* exposed to heat shock at 35°C for 1 hour and recovery at 20 °C for 5 hours. Each group contains at least 60 worms. **** P-val <0.001.

*C. elegans* adults exhibit a time-dependent decline in stress responsiveness coinciding with the onset of egg laying, including reduced inducibility of heat shock proteins^42^. Contrary to the substantial decrease in the inducibility of *mCherry::hsp-16*.*41* signal in aging wild type animals (**Figure 3B**), *znf-236* mutants displayed similar *mCherry::hsp-16*.*41* levels in the progression towards egg laying onset under no heat shock conditions (**Figure 3B, left**). This result indicates that ZNF-236 loss induces iHSPs in an age-independent manner, unlike the canonical heat shock response, whose strength declines with age.

We then examined heat inducibility in mutant animals as they age. In 1-day-old adult animals we observed allele-specific differences, with each reaching a distinct average induction level and displaying varying degrees of inducibility (**Figure 3B, right)**. Despite these differences, all strains showed reduced inducibility at day 1 compared to young adults, indicating that the canonical heat shock pathway remains intact and age-sensitive in *znf-236* mutants.

### *hsp-16*.*41* in a non-native chromosomal context retains heat-inducibility but loses responsiveness to *znf-236* inactivation

Next, we asked whether the *hsp-16*.*41* gene—including its promoter, open reading frame (ORF), and 3′ untranslated region (3′UTR)—is sufficient to drive *hsp-16*.*41* induction in *znf-236* mutants. If this minimal *hsp-16*.*41* unit alone were sufficient for its induction in the absence of ZNF-236, it would support a model in which regulation—or the maintenance of *hsp-16*.*41* quiescence—occurs direct engagement of the *hsp-16*.*41* gene by ZNF-236. Conversely, if *hsp-16*.*41* is not induced when isolated from its native genomic locus, it would suggest that ZNF-236 regulates its expression via long-range or locus-specific chromosomal interactions.

The *hsp-16*.*41* promoter contains well-characterized regulatory elements, including two heat-shock elements (HSEs; nTTCnnGAAnnTTCn), which are the widely conserved HSF-1 binding motifs in all eukaryotes (**Figure 4A**). This promoter is sufficient both for maintaining transcriptional quiescence under non-stress conditions and for enabling robust induction of *hsp-16*.*41* expression upon heat shock^43^.

**Figure 4.**
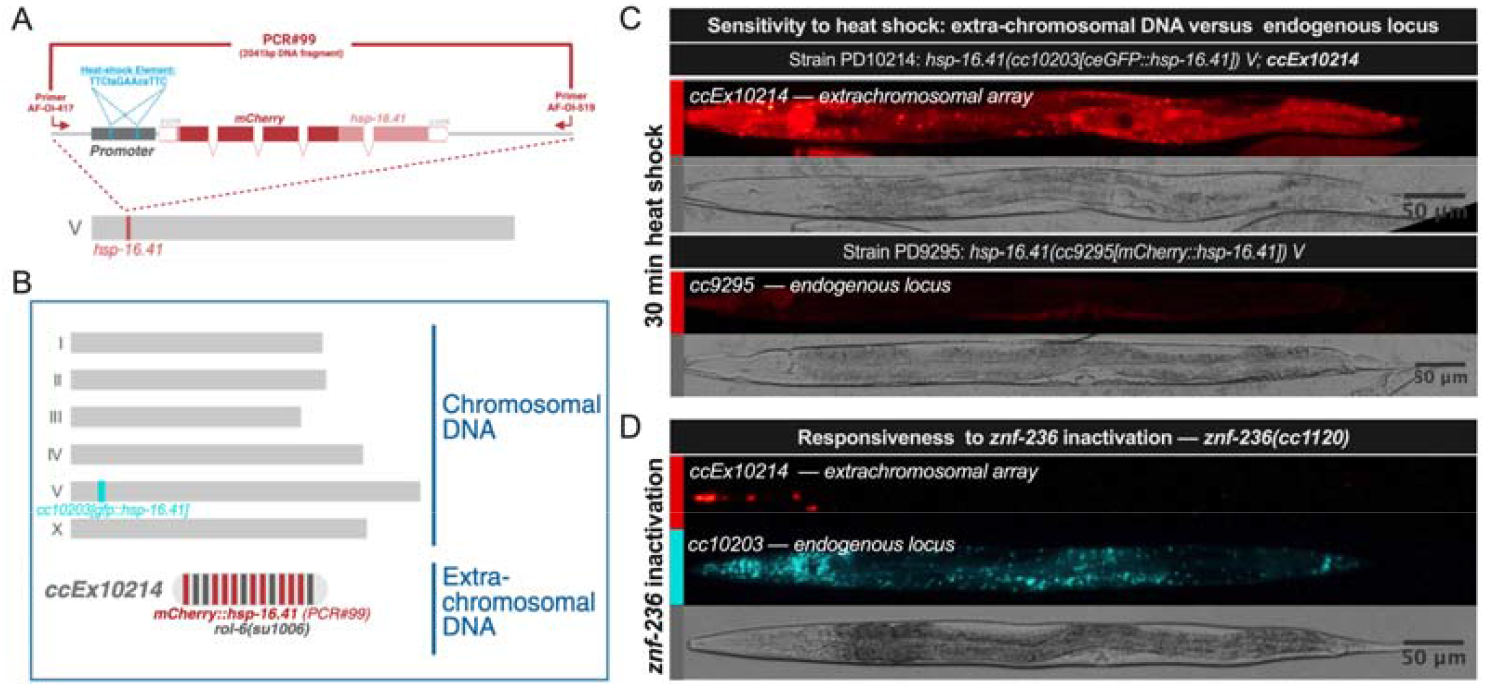
*hsp-16*.*41* in a non-native chromosomal context retains heat-inducibility but loses responsiveness to *znf-236* inactivation. (A) A ~2kb genomic fragment (termed PCR#99) containing the *mCherry::hsp-16*.*41* reporter and its native flanking sequences was PCR-amplified from the PD9295 reporter strain using a specific primer pair. (B) A stably inherited extrachromosomal array (*ccEx10214*) was generated by injecting PCR#99 along with a co-injection marker *rol-6(su1006)* into worms harboring an endogenously tagged *gfp::hsp-16*.*41* locus. (C) After a 30-minute of heat shock followed by 5-hour recovery, *mCherry::hsp-16*.*41* expression was higher in *ccEx10214* animals compared to animals carrying the single-copy endogenous reporter, indicating increased heat sensitivity of the multi-copy extrachromosomal array. (D) However, this multi-copy, hyper-heat-sensitive extrachromosomal *mCherryy::hsp-16*.*41* failed to respond to *znf-236* inactivation.

We first amplified and purified the genomic region spanning *hsp-16*.*41* gene using two PCR primers flanking the locus, with genomic DNA from the reporter strain PD9295 (**Figure 1A**) as the template (**Figure 4A**). This resulting PCR product (PCR#99) includes the *hsp-16*.*41* promoter, *mCherry*-fused ORF, and 3’UTR (**Figure 4A**). Then we injected this DNA fragment into the gonads of worms carrying a GFP-tagged *hsp-16*.*41* at its endogenous locus, which served as a reporter for *znf-236* activity. To identify transformants, we co-injected a linearized plasmid containing the dominant *rol-6(su1006)* mutation, a standard co-injection marker that causes a distinctive rolling phenotype. In *C. elegans*, injected DNA forms extrachromosomal arrays that can be stably inherited across generations (**Figure 4B**)^44^.

To test whether the *hsp-16*.*41* gene (PCR#99 DNA) responds appropriately to elevated temperatures, we heat-shocked the transgenic animals and confirmed the robust induction of the transgene (**Figure 4C**). Notably, the *mCherry::hsp-16*.*41* transgene on the extrachromosomal array appeared more sensitive to elevated temperatures than the endogenous *hsp-16*.*41* locus, likely due to the multi-copy nature of the array (**Figure 4C**). This result indicates that the *hsp-16*.*41* gene alone—independent of its native chromosomal context—is sufficient to respond to elevated temperatures. It is important to note that, this heat-induced activation likely reflects the accumulation of denatured proteins that activate HSF-1, which then can bind the heat-shock elements (HSEs) in the *hsp-16*.*41* promoter and activate it.

Next, to test whether the core *hsp-16*.*41* gene alone is regulated by ZNF-236, we introduced a *znf-236* mutation into the transgenic strain carrying the *hsp-16*.*41* extrachromosomal array. We found that the *hsp-16*.*41* expressed from the array (marked with *mCherry*) was not responsive to *znf-236* inactivation (**Figure 4D**). In contrast, as expected, the endogenous *hsp-16*.*41 locus* (marked with GFP) remained responsive (**Figure 4D**).

This result indicates that *hsp-16*.*41* induction in *znf-236* mutants requires regulatory context beyond those required for heat induction. This is not readily consistent with: 1) a model where *znf-236* inactivation leads to the accumulation of denatured proteins or general activation of HSF-1, which would be sufficient to induce *hsp-16*.*41* expression from extrachromosomal array DNA; 2) a model with universal sequence-specific binding of ZNF-236 the promoter of *hsp-16*.*41* to prevent an otherwise-default activation in the absence of stress. Instead, this finding supports a model in which ZNF-236 regulates gene expression through mechanisms dependent on chromosomal context--such as long-range enhancer-promoter interactions or broader features of chromatin organization. In other words, *hsp-16*.*41* regulation by ZNF-236 appears to depend on its native genomic environment or on chromatin features present in chromosomes but absent from extrachromosomal arrays.

### Genome-wide and chromosomal context-dependent activation of *hsp-16*.*41* in *znf-236* mutants

Next, we wanted to test whether ZNF-236 regulates *hsp-16*.*41* exclusively at its endogenous locus or can also regulate it at other genomic locations. To address this, we used miniMos transgenesis to randomly insert *mCherry::hsp-16-41* (PCR#110; almost identical to PCR#99 in Figure 4A) into the genome^45^ (**Figure 5A**). Using inverse PCR and Sanger sequencing of the insertion junctions, we mapped 23 independent insertion sites (**Figure 5A**)— all of which showed robust responsiveness to heat stress, as expected. This confirms that the core *hsp-16*.*41* promoter contains the necessary elements for both quiescence and inducibility, and that these elements are functionally robust regardless of chromosomal localization or context.

**Figure 5.**
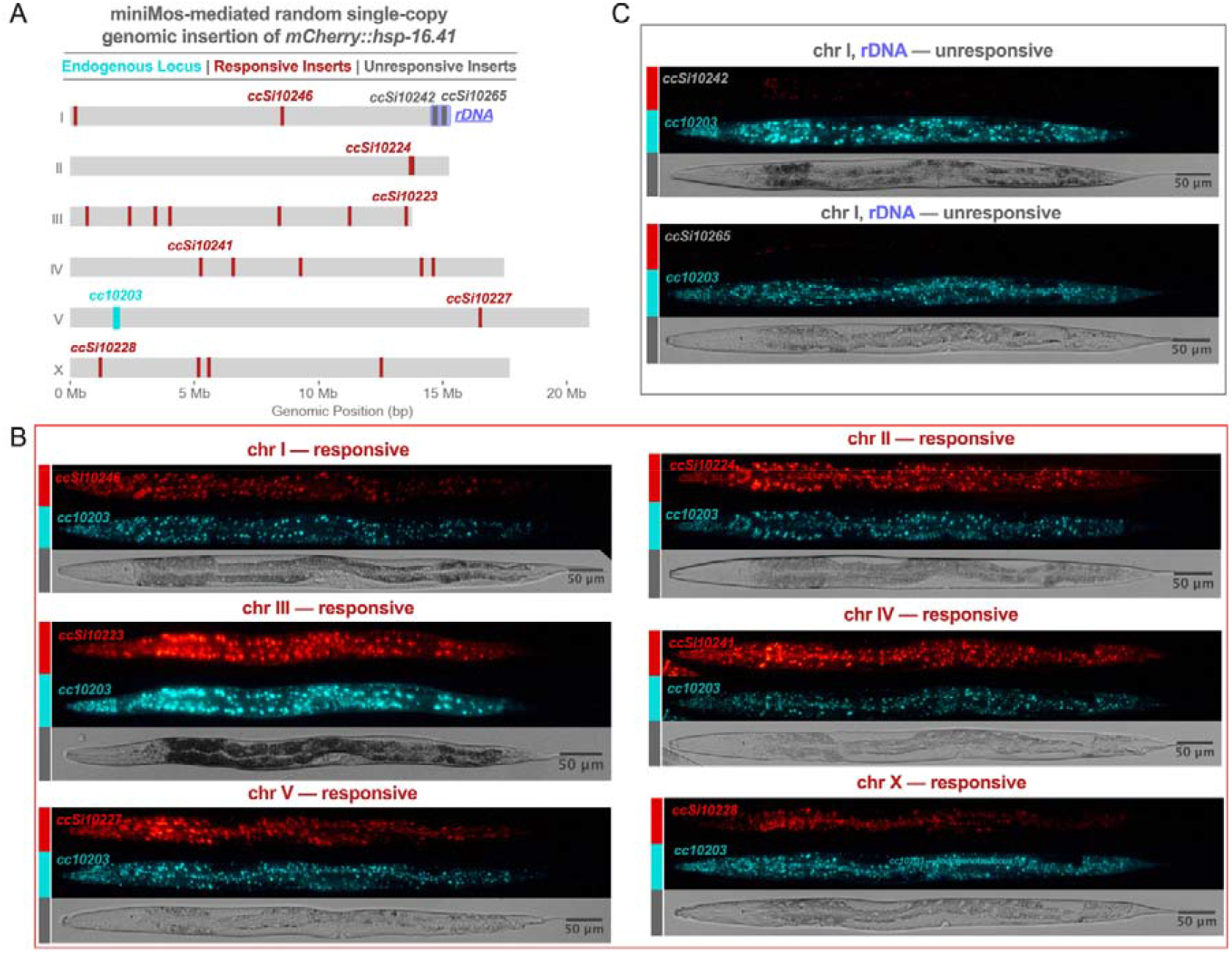
Genome-wide and chromosomal context-dependent activation of *hsp-16*.*41* in *znf-236* mutants. (A) The *mCherry::hsp-16*.*41* gene locus (Figure 4A) was inserted into various chromosomal locations using the MiniMos transposon-based single-copy insertion method. All resulting insertions were heat-inducible, confirming the functionality of the minimal *hsp-16*.*41* construct. 21 of 23 insertions also showed de-silencing in *znf-236* mutants, indicating genome-wide regulatory influence of ZNF-236. The two inserts that failed to respond to *znf-236* inactivation were both located within the ribosomal DNA (rDNA) repeat region at the distal end of chromosome I. (B) Representative micrographs showing induction of six (one per chromosome) MiniMos insertions in response to *znf-236* inactivation. In all six cases, *znf-236* is mutated (the allele *cc1147*) and the endogenous *hsp-16*.*41* is tagged with gfp (the allele *cc10203*). (C) Representative micrographs of the two rDNA-associated insertions that were unresponsive to *znf-236* inactivation are shown. Notably, the endogenous *hsp-16*.*41* locus (marked with gfp) remains responsive to znf-236 inactivation in these same animals, reinforcing that ZNF-236-mediated silencing is context-dependent—as observed with extrachromosomal DNA.

To our surprise—and inconsistent with a model involving a long-range cis-regulatory element acting specifically at the native locus—21 out of 23 insertion sites were also responsive to *znf-236* inactivation (**Figure 5A**). This shows that ZNF-236 can regulate gene expression at most loci across the genome, including at least one site on each chromosome (**Figure 5A and 5B**). These findings indicate that ZNF-236-mediated silencing is not restricted to the native *hsp-16*.*41* locus but instead suggests a role in a default silencing mechanism that operates broadly across most of the genome.

Notably, the two insertions that did not respond to *znf-236* inactivation—though still inducible by heat shock—were both located at the distal end of chromosome I, a region that harbors ribosomal DNA (rDNA) composed of tandem repeats encoding ribosomal RNA (**Figure 5A and 5C**). This provides further evidence that the *hsp-16*.*41* promoter alone, in the absence of appropriate chromosomal context, is not sufficient for regulation by ZNF-236.

Overall, these findings suggest that a ZNF-236-dependent default silencing mechanism restrains *hsp-16*.*41* expression across the genome. rDNA and extrachromosomal arrays may bypass this requirement—either because they lack the chromosomal context necessary to activate stress-induced genes in the absence of ZNF-236, or they engage alternative silencing mechanisms that function independently of ZNF-236.

## DISCUSSION

In this study, we leveraged the genetic tools in *C. elegans* to construct a sensitive and mutagenesis-compatible heat shock response (HSR) reporter by endogenously tagging *hsp-16*.*41*, a key inducible heat shock protein (iHSP) and a major HSF-1 target. We used this reporter to conduct a genetic screen to identify factors that maintain *hsp-16*.*41* expression quiescence under normal conditions. This screen identified that *znf-236*, which encodes the multi-zinc-finger protein ZNF-236 that is essential for suppressing *hsp-16*.*41* expression in the absence of stress. Further analysis showed that *hsf-1* is largely required for *hsp-16*.*41* induction in *znf-236* mutants, indicating that *znf-236* functions through or in parallel to *hsf-1* (**Figure 6**). Additionally, loss of *znf-236* led to the induction of not only *hsp-16*.*41* but also other iHSPs, known targets of HSF-1, and a subset of prion-like proteins previously described as stress regulated.

**Figure 6.**
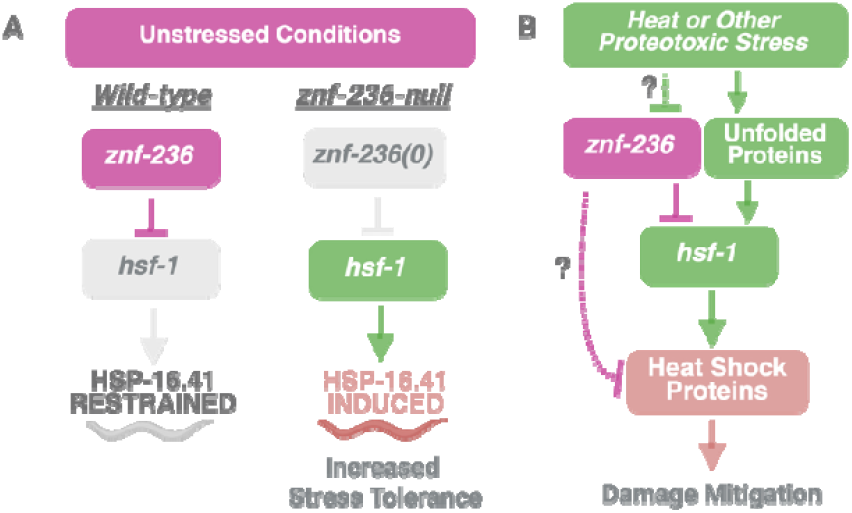
Observed genetic interactions and outcomes, with a model illustrating a potential role of ZNF-236 in the heat shock response pathway. (A) Under unstressed conditions, ZNF-236 restrains *hsp-16*.*41* expression. In the absence of *znf-236*, HSF-1-dependent *hsp-16*.*41* expression is observed. *znf-236* mutants also exhibit increased stress tolerance, the strength of which correlates with the level of *hsp-16*.*41* expression. (B) Models for ZNF-236 in the heat-shock pathway. ZNF-236 activity may be dampened by heat or other proteotoxic stress (green dotted line, question mark), leading to the expression of inducible heat shock proteins. This modulation could shape the rapidity, magnitude, or duration of the response, in parallel to the canonical unfolded protein/HSF-1 pathway. Alternatively, regulation of *znf-236* activity alone, without stress causing protein unfolding, could represent a distinct stress-response branch. The epistasis analysis (Figure 1F) uses a hypomorphic *hsf-1* allele, as null alleles are inviable. In worms containing both *znf-236* and *hsf-1* mutant alleles, *mCherry::hsp-16*.*41* expression is observed in some head tissues. This expression could result from residual HSF-1 activity or from HSF-1-independent activation of *hsp-16*.*41* in *znf-236* mutants. Therefore, we cannot rule out an hsf-1-independent regulation of hsp genes by *znf-236* (purple dotted, question mark).

To explore various cis- and trans-acting regulatory possibilities, we examined the positional requirements for functional regulation by ZNF-236. A simple model could be envisioned in which ZNF-236 is a trans-acting factor which might be directly responsible for *hsp* gene repression under nonstress conditions. However, our data support a model that challenges conventional expectations: ZNF-236 can silence targets irrespective of genomic position (arguing against certain cis-acting regulation models), yet its regulation is chromosomal-context dependent (arguing against the very simple model of direct ZNF-236 binding to *hsp-16*.*41* fully specified by sequences within this gene). In our working model, ZNF-236 is proposed to act through modification of local chromosome context, representing a novel regulatory mechanism for stress-responsive genes.

Loss of ZNF-236 responsiveness of the minimal *hsp-16*.*41* cassette on extrachromosomal arrays indicated that regulation by ZNF-236 requires additional cis-elements or chromosomal context absent from the array DNA. To examine functional requirements for regional chromosomal context and local sequence in ZNF-236-driven HSP regulation, we used a transposon-based insertion method that allowed insertion of an *mCherry-hsp-16*.*41* transgene at diverse genomic loci. The resulting set of random-site insertions of tagged HSP genes showed induction upon ZNF-236 loss at a wide variety of chromosomal sites. Only two exceptions to this induction were found, with both of these insertions present in the tandemly repeated ribosomal DNA (rDNA) loci in *C. elegans*. As an important positive control, we observed strong induction at all positions (including the rDNA repeat loci) following a heat shock. Thus, instead of finding a specific chromosomal region required for ZNF-236-mediated regulation, we discovered that ZNF-236 silences chromosomally encoded *hsp-16*.*41* irrespective of its genomic location, but in a manner sensitive to the chromosomal context. These findings revealed a previously unrecognized default silencing mechanism that restrains stress-inducible genes across most of the genome in *C. elegans*.

HSF-1 is largely required for *hsp-16*.*41* induction in *znf-236* mutants (**Figure 1F**); despite this, there are substantial differences observed between the responses to heat shock induction, known to occur through widespread protein denaturation, and *znf-236* loss. These differences include (1) gene expression changes in *znf-236* mutants overlap but are substantially distinct from those induced by heat shock (**Figure 2B**), arguing against *znf-236* being a component of canonical HSR activation, yet the subset of inducible chaperone genes is sufficient to confer thermotolerance (**Figure 3A**), suggesting sufficiency in mounting a biologically relevant heat shock response; (2) day 1 *znf-236* mutants maintain *hsp-16*.*41* induction despite age-related attenuation of the heat shock response (**Figure 3B**); (3) *hsp-16*.*41* encoded on extrachromosomal arrays or inserted into the rDNA locus is readily activated by heat stress— presumably via protein denaturation—but not by *znf-236* inactivation (**Figure 4**), suggesting that canonical heat-induced protein denaturation is unlikely the trigger for *hsp-16*.*41* activation in *znf-236* mutants. Together, these findings suggest that ZNF-236 may prevent activation of stress-induced genes not by modulating protein folding, but by surveillance of the chromosomal context or chromatin configuration that licenses HSF-1-mediated gene activation.

ZNF-236 and chromosomal context may regulate gene expression by restricting the access of activated HSF-1 to the *hsp-16*.*41* promoter or by modulating the activity of HSF-1 already bound at the promoter. HSF-1 is essential for *C. elegans* development—as well as for viability in yeast and proper development in mouse—and is known to occupy and regulate the promoters of numerous genes during this stage^23,46^. It is therefore plausible that this developmentally active, nuclear HSF-1—likely capable of inducing iHSPs—must be excluded from stress-responsive promoters under non-stress conditions. One potential role of ZNF-236 could be to maintain proper nuclear partitioning of active HSF-1. Accordingly, in *znf-236* mutants, this spatial regulation may be disrupted, allowing a portion of the active HSF-1 pool to access iHSP promoters, leading to their inappropriate activation.

The miniMos transgenesis --which we used to introduce a single-copy *hsp-16*.*41* gene into various loci in the genome (**Figure 5**)--clearly shows that a ZNF-236-mediated default silencing mechanism is capable of restraining *hsp-16*.*41* expression at most genomic loci tested. However, known gene regulatory mechanisms do not fully explain how this silencing can be mechanistically achieved. Notably, *hsp-16*.*41* itself, without the proper chromosomal context

--as exemplified by *hsp-16*.*41* in extrachromosomal DNA or inserted into the ribosomal DNA (rDNA) locus--is not regulated by *znf-236* (**Figure 4D and 5C**). It is therefore likely that iHSP de-silencing in the absence of ZNF-236 relies on features of chromatin architecture that are absent from extrachromosomal arrays or rDNA. It is also possible that extrachromosomal array DNA and rDNA do possess the potential to activate iHSPs, but they engage compensatory silencing mechanisms—such as a repetitive DNA silencing pathway^47–49^--suppressing iHSPs even in the absence of ZNF-236.

Several models --that are not mutually exclusive--could explain the observations in this study. In the first model, ZNF-236 activates a genetic target that is needed for suppression of stress response or suppresses a target sufficient to activate the response. While no obvious targets of either class emerge in comparing differential regulation in mutants, such models can certainly not be ruled out.

In a second model, distal enhancers distributed across the genome—at least one or more enhancer per chromosome—with the power to activate the *hsp-16*.*41* promoter could explain how ZNF-236 can regulate stress-induced genes across the genome. In this scenario, ZNF-236 or the default silencing pathway prevents chromosomally encoded *hsp-16*.*41* from interacting with this set of enhancers.

A third model posits the existence of ‘iHSP-activating’ nuclear compartments—such as the A and B (also called euchromatin and heterochromatin), active transcriptional hubs, or phase separated microenvironments^50^—where multiple active promoters, core transcriptional machinery (e.g. the RNA Pol II complex), and regulatory transcription factors like HSF-1, are concentrated. These iHSP-activating nuclear compartments are accessible by most chromosomal regions--except for the rDNA arm of chromosome I, which is likely sequestered in the nucleolus. In this model, ZNF-236 either prevents *hsp-16*.*41* from accessing these transcriptionally active compartments or modulates the activity of the compartment activity itself, for example by regulating the partitioning of HSF-1 to these domains.

ZNF-236 belongs to a large family of C2H2 multi-zinc-finger proteins in *C. elegans*, which includes over a hundred members^32^. This class of zinc-finger proteins is also the most abundant group of transcription factors in humans, where nearly half of them contain an effector domain known as KRAB (Krüppel-associated box)^32^, which recruits the transcriptional co-repressor KAP1^51^. Through this interaction, KRAB-ZNF proteins direct the silencing of specific genomic regions, often those harboring transposable elements^52^. Moreover, the KRAB-ZNF-KAP1 system can mediate long-range gene regulation through spreading of chromatin features^53^. A similar mechanism could explain how ZNF-236 might regulate gene expression from a distance. Importantly, the KRAB domain is found only in tetrapods and is absent in ZNF-236. Nevertheless, it remains possible that, despite lacking a KRAB domain, ZNF-236 recruits an analogous corepressor or other chromatin-modifying factors to mediate transcriptional repression at its target loci from a distance.

Some other mammalian multi-C2H2 zinc-finger proteins that lack effector domains, such as CTCF and YY1, regulate gene expression by shaping the three-dimensional structure of chromatin^54,55^. Both CTCF and YY1 can self-dimerize and promote chromatin looping, bringing distant regions of DNA into close spatial proximity. CTCF-mediated loops help define topologically associated domains (TADs), whereas YY1-mediated loops bring enhancers and promoters together^54,55^. The domain architecture of ZNF-236 closely resembles that of CTCF and, to a lesser extent, YY1—featuring a cluster of C2H2 zinc-finger domains interspersed with intrinsically disordered regions. This structural similarity --along with the findings in this study--raises the possibility that ZNF-236 may also function in organizing chromatin architecture. Accordingly, loss of ZNF-236 could lead to aberrant chromatin looping, resulting in ectopic activation or repression of genes affected by altered spatial interactions in the genome.

The presence of a default silencing mechanism controlling stress-induced genes, as described in this study, has several important implications. First modulating ZNF-236 activity— effectively enacting a regulated de-silencing—could enable a stress-responsive transcriptional program without fully activating the heat shock response **(Figure 6)**. Such a regulated de-silencing program, triggered by ZNF-236 inactivation might not require protein denaturation for its induction (**Figure 6B**), yet could still confer benefits under various stress conditions as we have observed in our thermotolerance assays and retained expression levels during aging (**Figure 3**). Second, ZNF-236 inactivation itself might be an integral component of the heat shock response: heat stress not only activates HSF-1 through protein denaturation but may also inactivate ZNF-236— potentially regulating the rapidity, magnitude, or duration of the response (**Figure 6B**). This is supported by the observation that *znf-236* mutants retain *hsp-16*.*41* inducibility upon heat shock (**Figure 2E, 3B**). Lastly, this silencing mechanism could help maintain the quiescence of stress-induced genes when they are relocated to new genomic contexts, such as through the activity of mobile genetic elements.

Together, our findings position ZNF-236 as a novel regulator that safeguards the quiescence of stress-induced genes in *C. elegans* likely by modulating chromosomal or nuclear context. Beyond classical models of stress response regulation --which emphasize transient activation of transcription factors by stress signals--this screen and observations with *znf-236* reveal a constitutive restraint of heat-shock genes by a default silencing mechanism likely essential for a well-tuned stress response. The evident role of a multi zinc-finger protein in this restraint provides a window into such silencing mechanisms, which may serve in a wide variety of physiological and pathological defense mechanisms.

## ACKNOWLEDGEMENTS

During the preparation of this work the authors used ChatGPT (GPT-4), a large language model, under human supervision, to edit some portions of the text and to write the Python scripts used for DEseq2 or TFD-Seq data analysis. The authors reviewed and edited the content as needed and take full responsibility for the content of the published article.

We thank Victor Ambros (UMass Chan Medical School), and Fire lab (Stanford University) members Karen Artiles and Dae-Eun Jeong for critical reading of the manuscript and helpful suggestions.

Funding: A.F. and O.I. are supported by NIH GM-R35-130366; R.M. is supported by NIH AG049665 and HF-GRO-23-11991705; A.G. is supported by NIH 5K00AG068511; D.J. is supported by R01AG086495.

## Materials and Methods

### *C. elegans* strains and Maintenance

All C. elegans strains were maintained under standard laboratory conditions on nematode growth medium (NGM) agar plates seeded with Escherichia coli OP50 as a food source. All strains used in this study are listed in **Supplemental Table S1**.

### CRISPR/Cas9-mediated tagging of *hsp-16*.*41*

To generate *hsp-16*.*41* reporter strains, CRISPR/Cas9-mediated genome editing was performed in *C. elegans* (N2, strain PD1099). Gravid adults were injected with a mixture of pre-assembled gRNA-Cas9 ribonucleoproteins (RNPs) targeting both *hsp-16*.*41* and the co-conversion locus *dpy-10*^56^, along with homologous recombination (HR) templates for fluorescent tagging.

#### gRNA assembly

A guide RNA targeting *hsp-16*.*41* (crRNA: AF-OI-351 5′-TTGAATCAGAATATGGAGAA-3′) was ordered from Integrated DNA Technologies (IDT) and annealed with tracrRNA (IDT) to form the functional gRNA. A second gRNA targeting the co-conversion marker *dpy-10* was prepared similarly. These two gRNAs were combined with Alt-R™ S.p. Cas9 Nuclease V3 (IDT) and incubated at room temperature to form RNP complexes.

#### Homologous recombination (HR) templates

The HR template for mCherry tagging (OI_CrisprMix#133) was PCR amplified (Thermo Fisher Scientific, Phusion High Fidelity DNA Polymerase Master Mix, cat#F531L) from plasmid pCFJ104 (Addgene #19328) using primers AF-OI-347 and AF-OI-356. The ceGFP HR template was PCR amplified from plasmid pCFJ2249 (Addgene #159847) using primers AF-OI-547 and AF-OI-548. PCR reactions (120 µL each) were purified using NucleoSpin® Gel and PCR Clean-Up kit (Takara Bio Usa, Inc. cat#740609) and added to the injection mixes at a final concentration of 160 ng/µL (mCherry) or 75 ng/µL (ceGFP).

Injected animals were recovered, and their progeny were screened for both the *dpy-10* co-conversion phenotype and fluorescence signal. F1 individuals expressing *mCherry* or *ceGFP* upon heat-shock were isolated, propagated, and genotyped by PCR using primers AF-OI-349 and AF-OI-350, followed by Sanger Sequencing to confirm precise insertions.

### CRISPR/Cas9-mediated mutagenesis of *znf-236*

To generate the *znf-236(cc1120)* deletion allele, two CRISPR guides targeting the 5′ and 3′ ends of the *znf-236* open reading frame (ORF) were co-injected. To express the guide RNAs from a plasmid, DNA primers (AF-OI-447/448, and AF-OI-449/450) were ordered from IDT, annealed, and cloned into pOI83 (a derivative of pRB1017 ^56^), yielding plasmids pOI354 and pOI355. These plasmids were then co-injected with a Cas9 expressing plasmid (pOI90) and a co-CRISPR plasmid targeting *unc-22* (pOI91) into strain PD9295.

Injected animals were recovered, and their progeny were screened for both the twitching phenotype and the constitutive (without external stress) expression the *mCherry::hsp-16*.*41*.

The *cc1120* deletion allele was identified by PCR using primers AF-OI-451 and AF-OI-372, followed by Sanger Sequencing.

The *cc9882* and *cc10213* alleles were isolated from the progeny of worms injected with a single guide RNA AF-OI-379 (IDT; 5′-GTATGCGCAACAGGTCTCGG-3′) targeting C-terminal sequences of *znf-236*. Injection mixes *OI_CrisprMix#148* and *OI_CrisprMix#186* were used to generate *cc9882* and for *cc10213*, respectively. Strain PD9295 was injected, and progeny were screened for *mCherry::hsp-16*.*41* induction in the absence of stress.

The *cc9882* allele was identified by PCR using primers AF-OI-371 and AF-OI-372, while *cc10213* was detected using primers AF-OI-408 and AF-OI-372. In both cases, Sanger Sequencing was used to confirm the identity of deleted sequences.

### Microscopy and Imaging

All DIC and fluorescent images were obtained using a Nikon Eclipse E600 equipped with an OMAX A3550U3 5.1 MP camera and ToupLite software (version OSX_19-05) software.

Before imaging, L4 stage animals (unless specified otherwise), cultured on standard NGM plates seeded with *E. Coli* OP50, were immobilized in 1 mM levamisole dissolved in M9 buffer. Images in Figure 1 as well as for 4 were captured using identical microscope settings, camera parameters, and exposure times across comparison sets (e.g. Figure 4D). Images were saved as TIFF files and processed uniformly using ImageJ (https://imagej.net/ij/); processing included selection, straightening, and alignment.

### RNA Sequencing and Transcriptome Analysis

Total RNA was extracted from two biological replicates of PD9295 (wild-type except for a *mCherry*-tagged *hsp-16*.*41* locus) and two biological replicates of *znf-236(cc1120)* mutant animals. Worms were collected at the L4 stage under non-stress conditions.

For each sample, 12 individual worms were picked into 20 µL of M9 buffer, followed by the addition of 300 µL of TRIzol reagent (Invitrogen). Samples were stored at −80°C overnight. The next day, all four samples were thawed and vortexed for 5 minutes at 4°C. 150 µL chloroform then added to each sample, mixed by inversion for approximately 15 seconds, and transferred to Phase lock gel Heavy tubes (QuantaBio, ref#2302830).

Following centrifugation at 10,000xg for 10 minutes at 4°C, the aqueous (top) phase was transferred to a new tube and cleaned using RNA clean & Concentrator™-5 kit (Zymo Research, cat#R1013) according to the manufacturer’s protocol.

#### rRNA depletion

To deplete ribosomal RNA (rRNA) prior to sequencing, 10 ng of total RNA was mixed with 25 ng of rRNA complementary DNA oligos (a pool of 94 DNA oligos AF-NJ-16 through AF-NJ-107, AF-NJ-150 and 151 purchased from IDT), and 2 µL of 5x hybridization buffer (500 mM Tris-HCl pH 7.4, 1M NaCl), in a total reaction volume of 8 µL.

Samples were denatured at 98°C for 2 minutes and annealed by slowly (0.1°C/sec) cooling down to 45°C. 2 µL of preheated (to 45°C) RNase H reaction mix was added (1 µL of Thermostable RNase H [5 units], 1 µL 10x digestion buffer: 500mM Tris-HCl, 1M NaCl, 200mM MgCl_2_). The reaction was incubated at 45°C for 1 hour, then cooled to 37°C.

To remove residual DNA, 3 µL of TURBO DNase (Thermo Fisher Scientific), 5 µL of TURBO DNase 10x buffer, 32 µL of nuclease-free water were added. The mixture was gently pipetted to mix and incubated for 30 minutes at 37°C. Following DNase digestion, 300 µL of STOP solution (1M ammonium acetate, 10mM EDTA, 0.2% SDS) was added to terminate the reaction. RNA was subsequently purified using RNA clean & Concentrator™-5 kit (Zymo Research, cat#R1013) according to the manufacturer’s protocol.

#### Library preparation, sequencing, and data analysis

RNA sequencing libraries were prepared using SMARTer Stranded Kit (Clontech Laboratories) following the manufacturer’s protocol. Libraries were sequences on an Illumina MiSeq platform. Adaptor trimming was performed with Trim Galore! (Version 0.6.7+galaxy0). Reads were aligned to the *C. elegans* genome (WS220) using HISAT2 (Version 2.2.1+galaxy1) and read counts were quantified using HTSeq-count (Version 0.9.1+galaxy1). Differential expression analysis was carried out using DEseq2 (Version 2.11.40.7+galaxy2). Volcano plots shown in Figure 2 were generated in Python using the Matplotlib library.

### MiniMos transgenesis

To generate single-copy insertions of *hsp-16*.*41*, we used MiniMos transposon-based integration system as previously described^45^. A ~2kb fragment (termed PCR#110) containing the *mCherry::hsp-16*.*41* open reading frame along with the native upstream and downstream sequences was PCR-amplified from the PD9295 strain using primers AF-OI-549 and AF-OI-558. This fragment was cloned into the MiniMos vector pCFJ1202, resulting in the final plasmid pOI383.

The pOI383 plasmid, together with pCFJ601 (Addgene #34874) and pCFJ420 (Addgene # 34877), was injected into *C. elegans* strain PD10203, which carries the *ceGFP*-tagged *hsp-16*.*41* at the endogenous locus.

Transgenic progeny were initially selected based on resistance to G418 (GoldBio), conferred by the NeoR cassette present in the pCFJ1202-based plasmid pOI383^45^. To eliminate individual worms carrying extrachromosomal arrays and enrich for single-copy integrants, heat shock was applied to induce the *peel-1* expression, which is encoded on the injected plasmid pOI383 but not retained following Mos1-mediated single-copy integration. Putative integrants were subsequently screened, and integration sites were identified using inverse PCR as previously described^45^.

### High throughput imaging

Worms were age-synchronized by bleaching. Briefly, animals are rinsed off the plate with M9 buffer and then centrifuged at 100 × g for 1 min. Worm pellet is then resuspended with a hypochlorite solution until adult worms are disintegrated. Eggs are then washed three times with M9 and incubated in an orbital shaker at 20 °C for 16 hours. The next day, the number of L1 worms was counted, and 300 were plated onto 6-cm NGM plates seeded with OP50. Worms were allowed to develop until the day 1 stage. We considered young adults as worms with an open vulva that have not yet started laying eggs. Day 1, animals indicate when they start laying eggs. Worms were heat shocked at 35 °C for 1 hour in a water bath with plates wrapped in parafilm. After the heat shock, worms were allowed to recover at 20 °C for 5 hours. To prepare worms for imaging, the plates were rinsed with M9 buffer, and the worms were allowed to settle in a conical 15-ml tube. After a couple of washes with M9 buffer, worms were resuspended with M9 buffer supplemented with 40mM sodium azide. A volume of 100 uL containing ~15-20 worms was dispensed into a well of 384-well plate (Greiner, 781091). The plate was imaged using an ImageXpress Micro confocal system. mCherry signal was collected using a filter with excitation (547nm) and emission 624/40nm with an exposure of 200 ms. A bright-field image was acquired with an exposure of 10 ms. Images were exported as TIFF files. A custom-trained Cellpose model^57^ was used to segment *C. elegans* bodies from the bright field image. The exported regions of interest (ROI) were used in a custom FIJI script to measure the *mCherry* intensity per worm.

### Thermotolerance assay

Strains were maintained at 20 °C. 10-15 gravid adults were transferred to an NGM plate seeded with OP50 and allowed to lay eggs for 2 hours. Synchronized worms derived from the eggs were allowed to develop until day 1 of adulthood. Twenty-five animals were transferred to four 35mm NGM plates, each seeded with OP50 per strain. Plates were then wrapped in parafilm and placed in a water bath set at 34 °C for 6 hours. 24 hours after recovering at 20 °C, animals were scored for movement. Dead animals were scored when they failed to move in response to a gentle touch.

