## Supplemental Figures for "A default silencing mechanism restrains stress-induced genes in *C. elegans*"

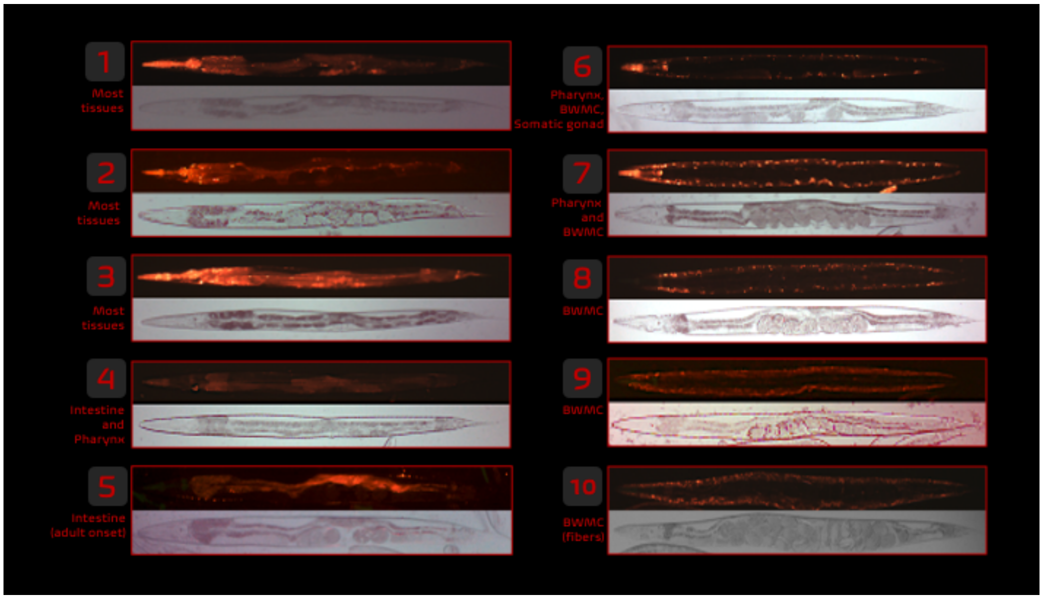


**Supplemental Figure 1. Mutagenized worm lines exhibiting constitutive *mCherry::hsp-16.41* expression in the absence of stress.**

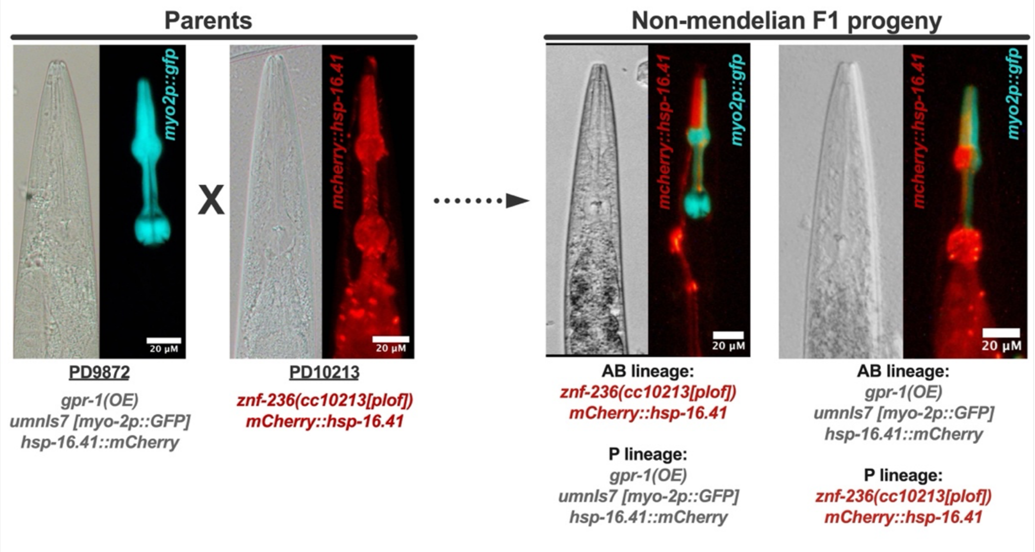


**Supplemental Figure 2. Cell-autonomous regulation of *hsp-16.41* by ZNF-236.** Site of action was probed by constructing a set of mosaic animals with *znf-236* loss-of-function in a defined subset of cells (cells derived from the embryonic P1 blastomere or the embryonic AB blastomere). An *mCherry* reporter present in all cells of these animals reports whether the stress induction is cell autonomous or non-autonomous. As pioneered by Besseling and Bringmann [33], motor protein GPR-1 overexpression allows separation of maternal and paternal genomes into AB and P daughter cells at the first cell division, creating the chimeric progeny with known lineages. To generate the animals shown, males carrying a *znf-236* mutation (strain PD10213) were crossed with *gpr-1*-overexpressing worms (strain PD9872), with both parents carrying the *mCherry::hsp-16.*41 reporter. In chimeric offspring, *mCherry::hsp-16.41* expression is observed only in cells that inherited the *znf-236* mutation, demonstrating cell-autonomous function of ZNF-236 in *hsp* gene silencing.
